# Ecological Interactions Drive Metabolomic Diversification in Amazonian *Pseudonocardia* Symbionts

**DOI:** 10.64898/2026.01.05.697768

**Authors:** Carlismari O. Grundmann, Weilan G. P. Melo, Andrés M. Caraballo-Rodríguez, Nina U. de A. Guardia, Ivan L. F. Migliorini, Ricardo R. Da Silva, Pieter C. Dorrestein, Norberto P. Lopes, Cameron R. Currie, Jon Clardy, Mônica T. Pupo

## Abstract

Fungus-growing ants engage in a multipartite symbiosis, including *Pseudonocardia* bacteria that produce antifungal metabolites to protect their fungal cultivar from the specialized pathogen *Escovopsis*. While different bioactive metabolites have been reported from ant-associated *Pseudonocardia*, most studies have focused on a limited number of strains, leaving the extent of chemical diversity across broader ecological contexts less resolved. Here, we investigated the antagonistic potential and metabolomic repertoires of 36 *Pseudonocardia* strains isolated from Amazonian *Paratrachymyrmex* ants. Pairwise bioactivity assays against two *Escovopsis* isolates revealed striking variability, with inhibition generally stronger and more diverse against the pathogenic fungus originating from the same ant genus. Untargeted LC-MS/MS metabolomics coupled with 16S rRNA-based phylogenetic analyses showed that closely related strains harbored highly divergent chemical profiles, underscoring a decoupling between taxonomy and metabolite output. Detailed analyses of selected isolates revealed the production of structurally diverse metabolites, including dentigerumycin analogs, provipeptide A, β-carbolines, and tetracycline-related compounds. Co-culture analysis uncovered metabolites absent in monocultures, including lichenysins, pepstatins, and hallobacillins, as well as conserved attinimicin, whose production was enhanced under pathogen challenge. These results highlight that both strain-specific metabolic repertoires and interaction-induced chemistry contribute to the defensive arsenal of *Pseudonocardia*. Together, our findings likely demonstrate that ecological pressures and local adaptation, rather than phylogeny alone, drive metabolomic diversification in this defensive symbiosis. Beyond their potential for novel bioactive compound discovery, these results provide insights into the chemical basis of multipartite symbioses, the dynamics of defensive mutualisms, and the ecological forces shaping microbial diversity in underexplored environments such as the Amazon.

**IMPORTANCE:** Microbial symbionts are central to host defense and natural product discovery, yet the factors driving their chemical diversification remain unclear. The fungus-growing ant–*Pseudonocardia*–*Escovopsis* system offers a powerful model to study how ecological context shapes microbial metabolism. By systematically characterizing multiple Amazonian *Pseudonocardia* strains, we show that antagonistic capacity and metabolomic repertoires vary widely, even among strains with highly similar 16S rRNA gene sequences, revealing a pronounced discordance between 16S-based phylogenetic relatedness and specialized metabolite production. These findings highlight the likely importance of ecological pressures and local adaptation in shaping metabolomic output, emphasizing symbiotic actinobacteria as both key ecological players and promising sources of antifungal natural products.

## INTRODUCTION

Fungus-growing ants (subtribe *Attini*) engage in a highly specialized multipartite symbiosis that has evolved over millions of years (1, 2). This mutualistic system includes the ants, a cultivated basidiomycete fungus that serves as their primary food source, and a suite of microbial associates with distinct ecological roles (3–6). Among the most well-characterized of these partners are actinobacteria from the genus *Pseudonocardia*, which are vertically transmitted across generations and colonize the cuticle of worker ants (3, 7). These bacteria function as defensive symbionts, producing antifungal compounds that selectively inhibit pathogens, most notably *Escovopsis*, a specialized mycoparasite that infects and consumes the fungal cultivar (3, 8, 9). The *Attini–Pseudonocardia–Escovopsis* interaction exemplifies a co-evolved defense strategy within a complex ecological network, offering an exceptional model for studying microbial symbiosis, chemical ecology, and the evolution of interspecies cooperation (10, 11).

Extensive research on *Pseudonocardia* strains isolated from attine ants has revealed the production of structurally diverse natural products, many of which exhibit potent antifungal activity. Notable examples include dentigerumycin, a cyclic depsipeptide isolated from *Apterostigma dentigerum*-associated *Pseudonocardia* through bioassay-guided fractionation (12); its less active analogues, the gerumicins, derived from strains associated with *Apterostigma* spp. and *Trachymyrmex cornetzi* (13); and selvamicin, a polyene macrolide discovered via genome mining and heterologous expression, showing potent activity against *Candida albicans* (14). These compounds are believed to play critical roles in defending the fungal cultivar from specialized pathogens such as *Escovopsis*, reinforcing the ecological relevance of these symbionts. However, most studies have focused on a limited number of strains, often from Central American collections, and the extent of metabolic diversity across broader geographic contexts, particularly from underexplored regions like the Amazon, remains largely unknown.

Understanding this diversity requires investigating how environmental and ecological variables influence microbial metabolism. Mounting evidence suggests that microbial specialized metabolism is shaped by ecological and biogeographical factors (15–18). In insect-associated actinobacteria such as *Pseudonocardia*, environmental variation and host interactions have been proposed to influence the expression of biosynthetic gene clusters and the production of specialized metabolites (13, 19). Additionally, ecological interactions, particularly those involving fungal antagonists like *Escovopsis*, are known to induce metabolomic shifts that reveal otherwise silent or low-abundance compounds. Yet, few studies have directly examined the chemical output of *Pseudonocardia* during such interactions, and comprehensive metabolomic analyses under ecologically relevant and comparative conditions remain rare (20, 21). In particular, large-scale studies involving multiple *Pseudonocardia* strains exposed to pairwise co-culture with fungal pathogens are lacking, despite their importance for distinguishing conserved responses from strain-specific metabolic strategies. These knowledge gaps highlight the need to explore how interspecies interactions and geographic context collectively shape the metabolic potential of microbial symbionts and drive chemical diversification within defensive symbioses.

Addressing this need, recent efforts have begun to survey the metabolic capabilities of larger collections of *Pseudonocardia* strains across geographic regions. For example, it was shown that *Pseudonocardia* symbionts associated with fungus-growing ants can produce the conserved antifungal compound attinimicin, with biosynthesis reported across multiple regions of Brazil but notably absent in strains from Panama (22). Building on these insights, we found that 36 *Pseudonocardia* strains isolated from distinct Amazonian locations harbor a remarkably broad and dynamic chemical repertoire, extending beyond previously described conserved metabolites. Through mass spectrometry-based metabolomics and microbial co-culture assays with two *Escovopsis* strains, we observed the production of structurally diverse compounds, including dentigerumycins, β-carbolines, provipeptides, structurally related tetracyclines, lichenysins, and pepstatins. Some of these compounds were specifically induced during co-culture with *Escovopsis* strains, including attinimicin. These findings support the idea that *Pseudonocardia* symbionts have evolved a flexible and ecologically responsive chemical arsenal, shaped by local selective pressures and long-term co-evolution. Collectively, our results underscore the important role of microbial symbionts as adaptive chemical defenders in complex multipartite symbioses.

## RESULTS

### Antagonistic potential of *Pseudonocardia* against *Escovopsis*

To evaluate the inhibitory activity of *Pseudonocardia* strains against parasitic *Escovopsis* fungi, 72 pairwise antagonism assays were performed using 36 bacterial strains from *Paratrachymyrmex* ant colonies, each tested against two fungal strains isolated from different ant genera: ICBG 729 (*Paratrachymyrmex* colony) and ICBG 740 (*Acromyrmex* colony). Identification codes and the geographic coordinates within the Brazilian Amazon region for the *Pseudonocardia* and *Escovopsis* strains are provided in **Table S1**.

The inhibition profiles were scored on a scale from 0 to 4, based on qualitative assessment of fungal growth restriction, where level 0 shows complete fungal overgrowth, including over the bacterial streak; level 1 displays limited fungal restriction; level 2 reflects partial inhibition; level 3 demonstrates strong but incomplete suppression; and level 4 reveals full fungal growth inhibition with no visible colonization beyond the inoculation site. These results are summarized in a heatmap (**Figure 1A**).

**Figure 1.**
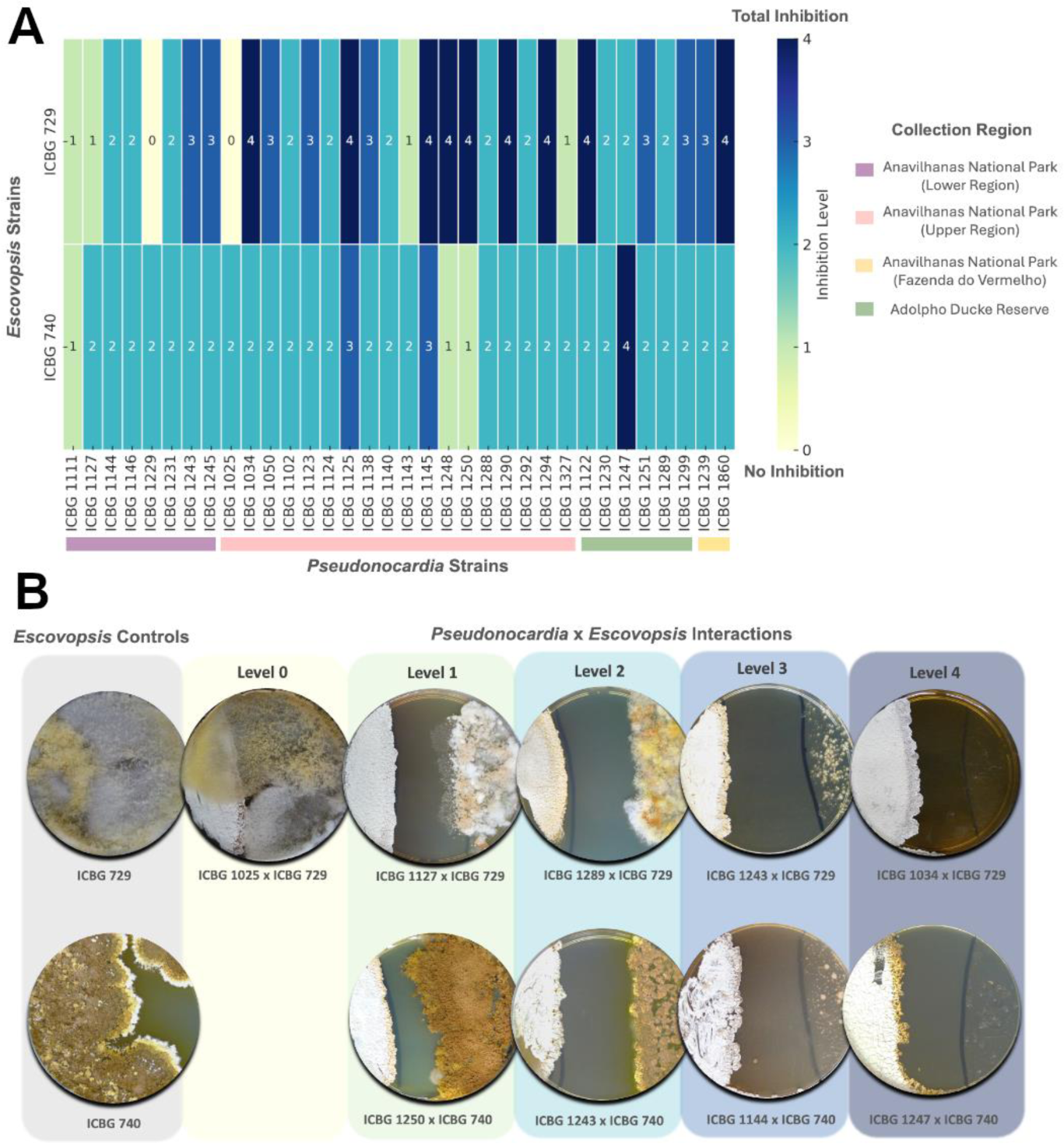
Pairwise interactions between *Pseudonocardia* and *Escovopsis* strains. **(A)** Heatmap showing inhibition levels of 36 *Pseudonocardia* strains, isolated from *Paratrachymyrmex* ant nests, against two different *Escovopsis* strains (ICBG 729 and ICBG 740). Each number represents a specific level of fungal inhibition, ranging from 0 (no inhibition, light yellow) to 4 (strong inhibition, dark blue), as indicated by the color gradient. *Pseudonocardia* strains are grouped by collection region, highlighted by colored bars at the bottom: Anavilhanas National Park – Lower Region (purple), Upper Region (pink), Fazenda do Vermelho (yellow), and Adolpho Ducke Reserve (green). *Escovopsis* strain ICBG 729 was isolated from a *Paratrachymyrmex* nest, while ICBG 740 was isolated from an *Acromyrmex* nest. **(B)** Representative in vitro antagonism assays between selected *Pseudonocardia* and *Escovopsis* strains after 14 days of co-culture, organized by inhibition level (0 to 4). On the left, monoculture controls of both *Escovopsis* strains. No antagonistic interactions with ICBG 740 exhibited level 0 inhibition.

Overall, a broad range of antagonistic profiles was observed, with several *Pseudonocardia* strains displaying strong inhibition (levels 3-4) against at least one *Escovopsis* strain. Notably, inhibition was generally stronger and more variable against ICBG 729, while responses to ICBG 740 were consistently weaker across most bacterial isolates, regardless of their region of origin. This contrast suggests that the identity of the fungal pathogen plays a key role in modulating the defensive response. Moreover, for ICBG 729, a pattern emerged in which highly inhibitory phenotypes were more frequently observed among *Pseudonocardia* strains from the Upper Anavilhanas region and “Fazenda do Vermelho”, hinting at the influence of local ecological pressures or co-evolutionary history. These results support the hypothesis that both pathogen specificity and geographic origin shape the antifungal potential of *Pseudonocardia* symbionts. Some representative co-culture phenotypes are shown in **Figure 1B**, illustrating the full spectrum of inhibition observed, from unrestricted fungal overgrowth (level 0) to complete fungal suppression (level 4) after 14 days of incubation.

### Metabolomic and phylogenetic profiles reveal strain-specific diversity

To further explore the molecular basis underlying the observed phenotypic diversity in antifungal activity, we investigated the metabolomic and phylogenetic profiles of *Pseudonocardia* strains. Given the wide range of inhibition patterns revealed in the antagonism assays, we hypothesized that these functional differences may be linked to variation in specialized metabolite production. Using liquid chromatography–tandem mass spectrometry (LC-MS/MS) based untargeted metabolomics and 16S rRNA gene phylogenetic analysis from *Pseudonocardia* strains, we assessed the chemical and evolutionary diversity across our bacterial collection. These analyses provide insight into whether metabolomic diversity reflects phylogenetic relatedness and to what extent metabolic potential aligns with antifungal performance.

The heatmap of detected ions (**Figure 2A**) reveals extensive chemical diversity across strains, with certain isolates displaying unique or highly enriched ion signatures (as ICBG 1122, ICBG 1025 and ICBG 1860), indicative of distinct specialized metabolite repertoires. Although *Pseudonocardia* strains are closely related taxonomically, reflected in the tight clustering of phylogenetic positions based on 16S rRNA sequences (**Figure S1**), many exhibit markedly divergent chemical profiles. Procrustes analysis (**Figure 2B**) (23) provides a comparative view of the relationship between phylogeny and metabolomic profiles across all strains by aligning two independent ordinations: one derived from a similarity matrix based on 16S rRNA gene sequences and the other derived from a metabolite feature abundance matrix. In this analysis, each strain is represented by a pair of nodes connected by two-segment colored edges, where one segment links the strain to its phylogenetic position (black segment) and the other links it to its corresponding metabolomic profile (gray segment). Short connecting bars and similar orientations indicate concordance between evolutionary relatedness and metabolite production, whereas long bars observed for many strains in our dataset highlight mismatches between phylogenetic similarity and chemical profiles.

**Figure 2.**
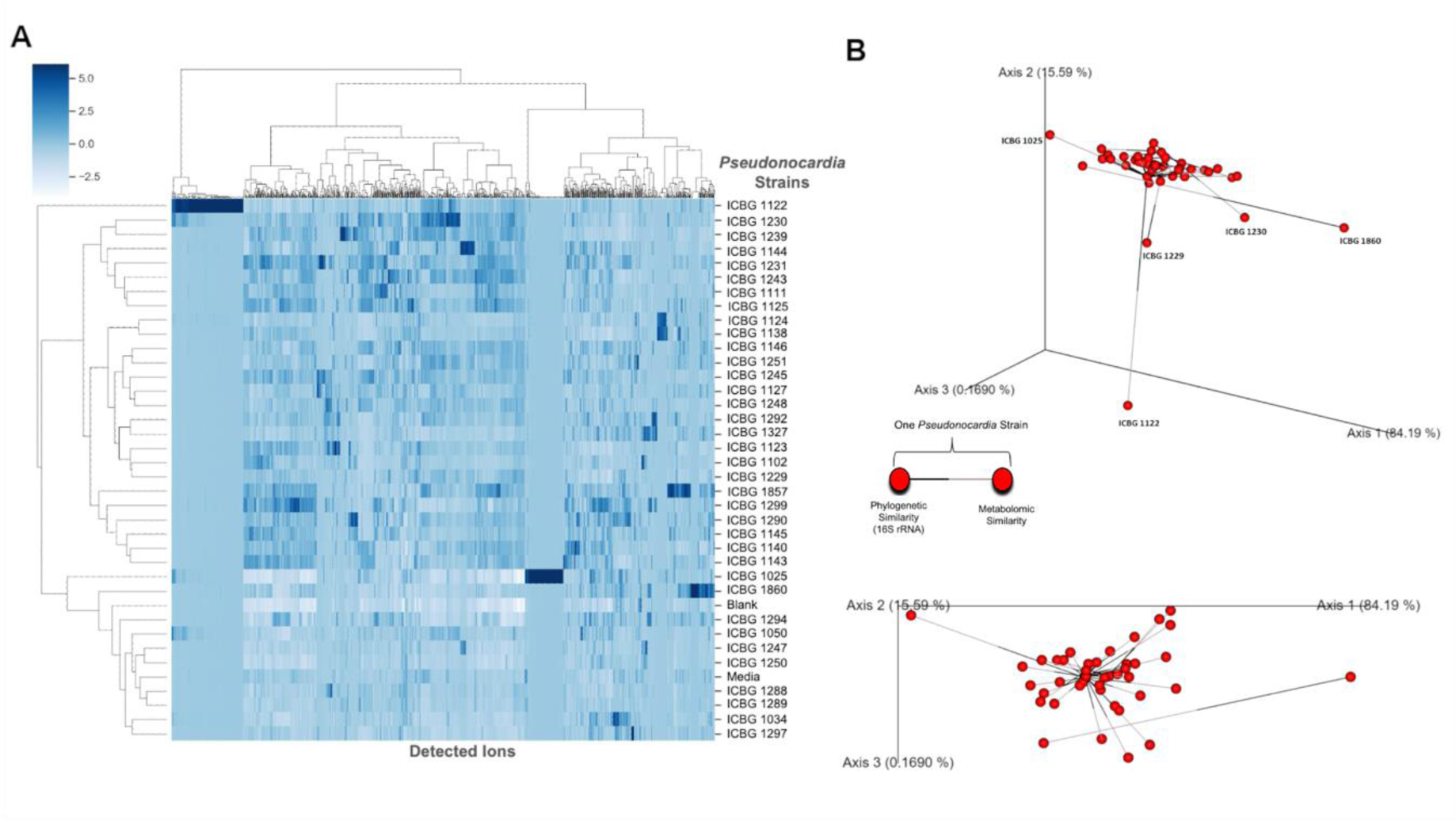
Metabolomic and phylogenetic analysis of *Pseudonocardia* strains. **(A)** Hierarchical clustering heatmap showing the relative abundance of detected ions in LC-MS/MS extracts from 36 *Pseudonocardia* strains. Each column represents a detected ion, and each row corresponds to a *Pseudonocardia* strain. Ion intensities are shown on a blue color scale, where darker shades indicate higher relative abundance. Clustering was performed on both axes to reveal patterns of metabolomic similarity among strains. **(B)** Procrustes analysis comparing phylogenetic relationships (based on 16S rRNA gene sequences, **Figure S1**) with metabolomic profiles (LC-MS data) of *Pseudonocardia* strains. Each strain is represented by a bar connecting two nodes: one for phylogenetic similarity (black ends) and one for metabolomic similarity (gray ends). The spatial alignment of bars indicates the degree of concordance between the two data modalities. Closer proximity of nodes across strains reflects similar phylogenetic and metabolomic profiles.

### Strain-specific specialized metabolite production and antimicrobial activity

Based on the observation above that metabolomic profiles vary substantially among closely related *Pseudonocardia* strains, we next focused on characterizing the specific metabolites responsible for these differences, particularly those enriched in strains with distinctive chemical and phenotypic profiles. We selected three strains (ICBG 1122, ICBG 1025, and ICBG 1860) based on their unique metabolomic signatures and divergent positions in the Procrustes analysis, which suggested they possess distinct specialized metabolic potential. Using molecular networking (24, 25) of LC-MS/MS data from *Pseudonocardia* monocultures and co-cultures extracts, combined with genome mining, and bioassays, we annotated several strain-specific metabolites with putative or confirmed antimicrobial activity. The following figures (**Fig. 3–5**) detail the chemical profiles of each strain, emphasize the most prominent compound families detected, and present in some cases the results of bioactivity assays that validate their functional relevance in inhibiting fungal and bacterial pathogens.

**Figure 3.**
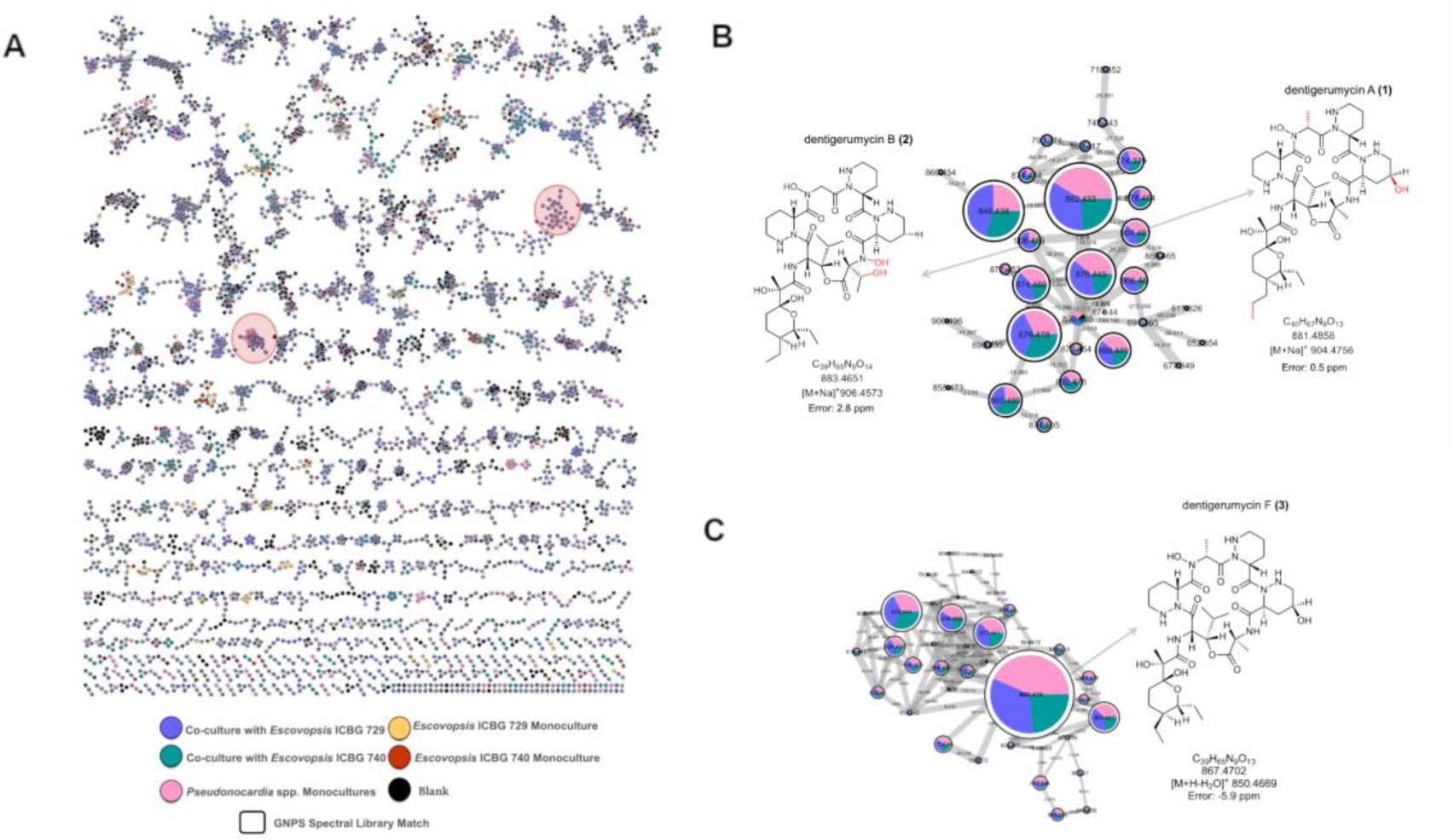
Molecular networks reveal dentigerumycin analogs produced by *Pseudonocardia* sp. ICBG 1122 and its interactions with *Escovopsis* spp. (A) Overview of the molecular networks constructed from LC-MS/MS data of 36 *Pseudonocardia* strains cultured in monoculture and in co-culture with *Escovopsis* ICBG 729 and ICBG 740. Nodes represent MS/MS spectra, with edges connecting structurally related molecules based on spectral similarity. Node colors correspond to sample types as indicated in the legend, node sizes reflect the number of MS/MS spectra (scans), and pie charts indicate the relative abundance of each feature in the different culture conditions. The two clusters highlighted with red circles correspond exclusively to metabolites from *Pseudonocardia* sp. ICBG 1122, either in monoculture or in co-culture. **(B)** Zoom-in of a molecular cluster derived from *Pseudonocardia* sp. ICBG 1122 and its co-cultures with *Escovopsis* spp., showing the distribution of ions and annotation of dentigerumycin A and dentigerumycin B as sodium adducts. **(C)** Zoom-in of another exclusive *Pseudonocardia* sp. ICBG 1122 cluster, showing annotation of dentigerumycin F as a protonated species. Self-loops are not shown.

**Figure 4.**
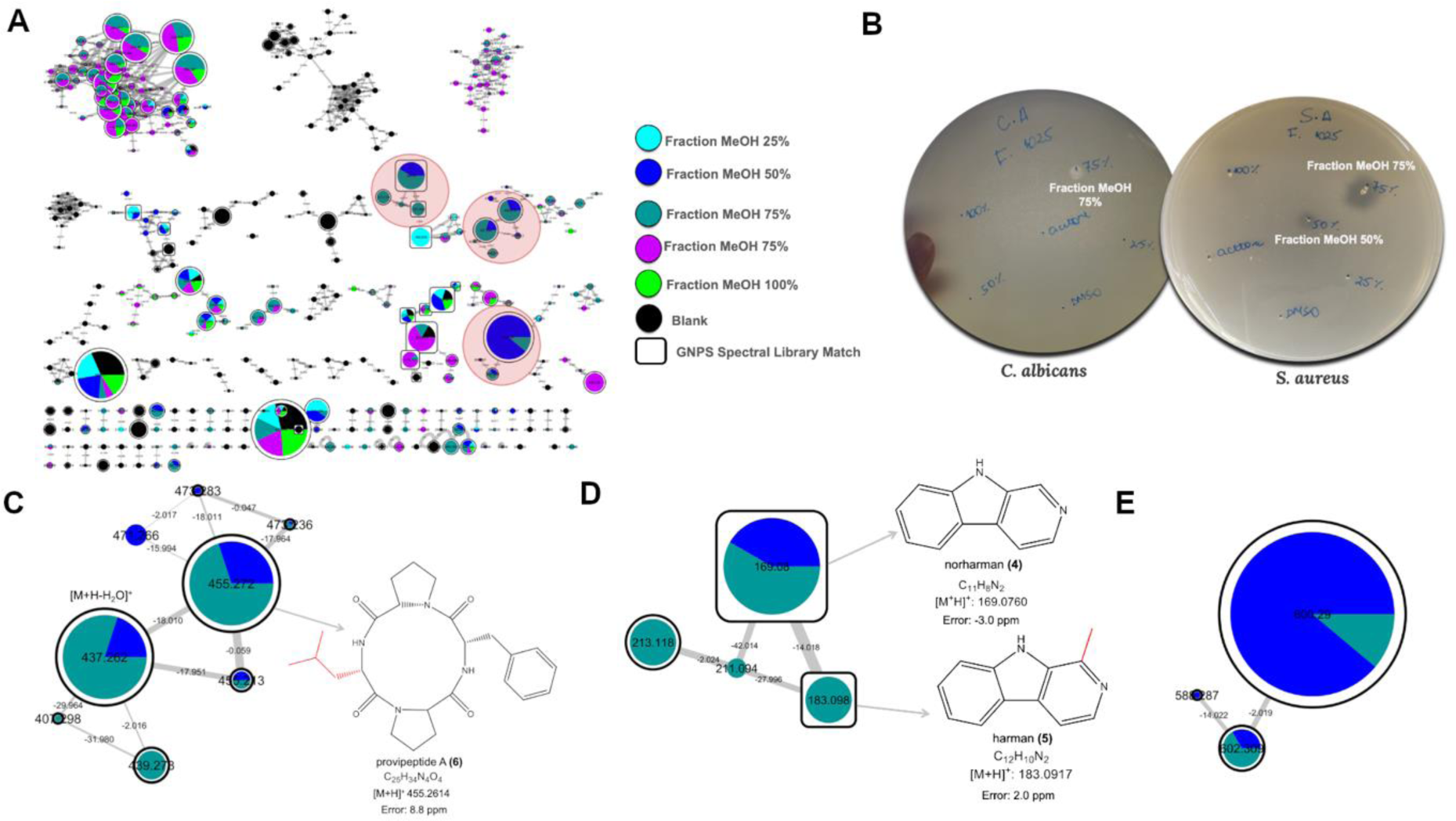
Molecular networks and antimicrobial activity of *Pseudonocardia* ICBG1025 extracts. **(A)** Molecular networks generated from LC-MS/MS data of methanol and SPE fractions obtained from small-scale fermentation of *Pseudonocardia* strain ICBG1025. Nodes represent MS/MS spectra, with edges connecting structurally related molecules based on spectral similarity. Node colors correspond to sample types as indicated in the legend, node sizes reflect the number of MS/MS spectra (scans), and pie charts indicate the relative abundance of each feature in the different methanolic fractions (25%, 50%, 75%, and 100%) and 100% acetone fraction. Black nodes represent ions detected in the blank control. Nodes outlined in white indicate GNPS library matches. Red circles highlight clusters enriched only with MeOH 50% and 75% fractions (blue and green, respectively). **(B)** Antimicrobial activity of the organic fractions (dissolved in DMSO at 10 mg/mL) against *C. albicans* and *S. aureus*. Clear inhibition zones are observed, particularly for the 50% and 75% MeOH fractions. **(C, D, E)** Selected molecular clusters predominantly found in the 50% and 75% MeOH fractions. **(C)** Cluster containing *m/z* 455.272, identified as the adduct [M+H]+ for provipeptide A with additional ions representing dehydrated and fragment ions. **(D)** Cluster annotated via GNPS as norharman (*m/z* 169.0760) and harman (*m/z* 183.0917 **(E)** Unannotated cluster with precursor ions *m/z* 502.309 and 600.297, found exclusively in the 50% and 75% MeOH fractions. Node size is proportional to the number of MS/MS scans acquired for each ion.

**Figure 5.**
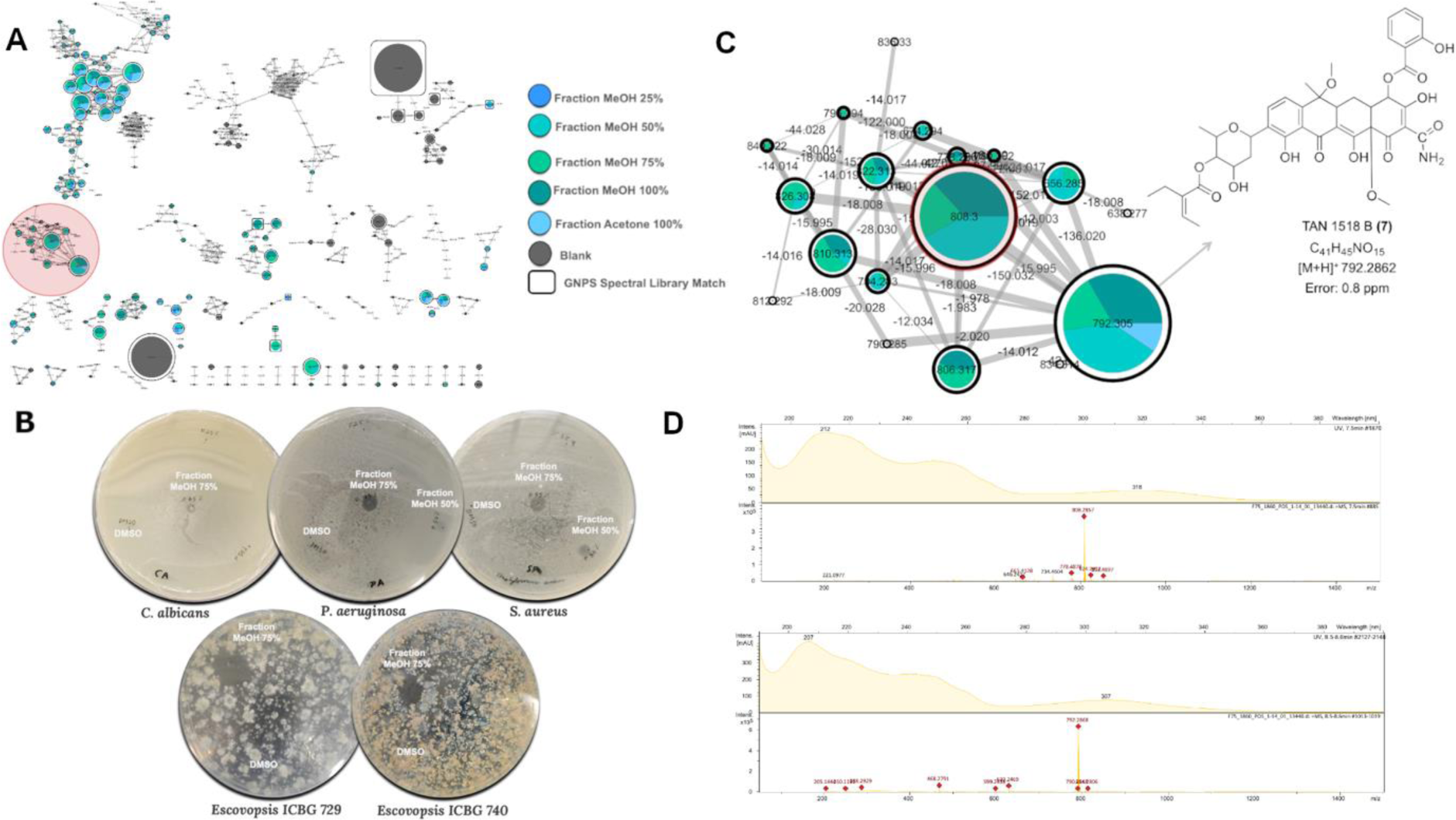
Molecular networks and antimicrobial activity of *Pseudonocardia* sp. ICBG 1860 extracts. **(A)** Overview of all ions detected in the LC-MS/MS analysis of small-scale cultures of *Pseudonocardia* sp. ICBG 1860, grouped by molecular similarity using GNPS molecular networking. Nodes represent MS/MS spectra, with edges connecting structurally related molecules based on spectral similarity. Node colors correspond to sample types as indicated in the legend, node sizes reflect the number of MS/MS spectra (scans), and pie charts indicate the relative abundance of each feature in the different methanolic fractions (25%, 50%, 75%, and 100%) and 100% acetone fraction. Gray nodes represent blank samples. A molecular cluster enriched in methanolic fractions 50%, 75% and 100% is highlighted in red. **(B)** Antimicrobial activity of MeOH/H₂O fractions of *Pseudonocardia* sp. ICBG 1860 extracts tested against *Candida albicans*, *Staphylococcus aureus*, *Pseudomonas aeruginosa*, and two *Escovopsis* strains (ICBG 729 and ICBG 740). Inhibition halos were recorded after 24 hours for human pathogens and after 14 days for *Escovopsis* strains. Notably, the 50% and 75% MeOH fractions showed significant bioactivity. **(C)** Detail of the molecular cluster highlighted in (A), corresponding to ions detected predominantly in the 50%, 75%, and 100% MeOH fractions. The ions at *m/z* 808.3 and *m/z* 792.3 are. The ion at m/z 792.3 was annotated as [M+H]^+^ of TAN 1518B. **(D)** UV-visible absorption spectra (measured in ACN at 254 nm) and MS1 spectra of the ions at *m/z* 808.3 and 792.3. The node sizes in the network are proportional to the number of MS/MS scans associated with each ion.

Strain ICBG 1122 selectively produces dentigerumycin analogs in response to *Escovopsis* challenge. Classical molecular networking analysis revealed *Pseudonocardia* sp. ICBG 1122 is chemically distinct, with two exclusive molecular clusters detected only in this strain under both monoculture and co-culture conditions (**Figure 3A**). These clusters, absent in all other strains, suggest a unique biosynthetic capacity.

Upon closer inspection, these two distinct molecular clusters revealed correspondence to different ionization forms of dentigerumycin-family compounds. The first cluster (**Figure 3B**) includes features annotated as sodiated adducts of dentigerumycin (*m/z* calc. 904.4756, *m/z* obsv. 904.4749, 0.5 ppm) and dentigerumycin B (*m/z* calc. 906.4573, *m/z* obsv. 906.4547, 2.8 ppm), based on accurate mass and low ppm error, consistent with MSI (Metabolomics Standard Initiative) level 3 identification (26)(**Figures S2**). The second cluster (**Figure 3C**) contains dentigerumycin F, detected as a protonated in source fragmented ion (*m/z* calc. 850.4669, *m/z* obsv. 850.4618, -5.9 ppm), which was isolated and structurally characterized based on spectrum comparison with literature, qualifying as MSI Level 1 (21)(**Figures S3-S4, Table S2**). Notably, dentigerumycin F was prioritized for isolation due to its higher production levels relative to other analogs. It is important to note that the separation of dentigerumycin-related features into two molecular clusters reflects differences in ionization state (sodiated versus protonated/in-source fragmented ions), which generate distinct MS/MS spectra and consequently lead to their segregation during molecular networking (27, 28).

Qualitative inspection of the pie charts within the molecular network (**Figures 3B–C**) suggests a slightly higher relative abundance of dentigerumycin analogs in co-culture with *Escovopsis* ICBG 729 compared to ICBG 740, indicating that metabolite induction may be strain-specific and context-dependent.

Additionally, genome analysis of strain ICBG 1122 revealed a biosynthetic gene cluster (BGC) with high similarity to known dentigerumycin pathways, reinforcing the annotation and underscoring the specialized metabolic potential of this symbiont in the defensive ant-microbe symbiosis (**Figure S5**).

Unlike strain ICBG 1122, *Pseudonocardia* sp. ICBG 1025 did not exhibit strong inhibition against *Escovopsis*, particularly strain ICBG 729. To investigate the molecular basis underlying its distinct metabolomic profile observed in the previous results, we performed a targeted chemical analysis, later tested against human pathogens. This approach aimed to correlate bioactivity with unique metabolite identities, likely related to the specific ion signatures shown in **Figure 2A**.

Small-scale monoculture cultivations were made, extracted and fractionated by solid-phase extraction (SPE) using a stepwise methanol (MeOH) gradient with water (25%, 50%, 75%, and 100% MeOH), followed by elution with 100% acetone. Aliquots from each fraction were analyzed by LC-MS/MS for molecular networking visualization (**Figure 4A**) and screened for antimicrobial activity against *Candida albicans* and *Staphylococcus aureus* (**Figure 4B**).

Molecular networking revealed three unique clusters present exclusively in the 50% and 75% MeOH fractions (highlighted in red, **Figure 4A**), which were also the most bioactive against the pathogens in the bioassays. The first cluster (**Figure 4C**) included the protonated ion related to provipeptide A (*m/z* calc. 455.2614, *m/z* obsv. 455.2654, 8.7 ppm) and one of its in-source fragmented ions (*m/z* 437.262 [M+H-H_2_O]^+^). This annotation was confirmed by isolation, NMR data and comparison with literature, being identified as MSI Level 1 (29)(**Figures S6-10**, **Table S3**). The second cluster (**Figure 4D**) was annotated via GNPS as protonated adducts, containing norharman (*m/z* calc. 169.0760, *m/z* obsv.169.0775, 3.0 ppm) and harman (*m/z* calc. 183.0917, *m/z* obsv. 183.0919, 2.0 ppm), corresponding to MSI Level 2 annotations based on spectral library matches (**Figures S11-12**). A third cluster (**Figure 4E**), containing ions at *m/z* 502.309 and 600.297, could not be annotated using the GNPS library or MS1 comparison with public natural products databases. Additionally, due to low abundance of these unannotated ions in the large-scale cultivation, isolation for NMR structural characterization was not possible (MSI Level 4).

In a similar way, the metabolomic profile investigation of *Pseudonocardia* sp. ICBG 1860 was conducted by doing small-scale cultivations followed by methanol-based solvent fractionation and LC-MS/MS analysis for molecular networking visualization (**Figure 5A**), in parallel with antimicrobial bioassays of the corresponding fractions (**Figure 5B**). In addition to the tested pathogens mentioned above, the fractions also showed activity against *Pseudomonas aeruginosa*, and *Escovopsis* ICBG 729. The 50% and especially the 75% MeOH fractions exhibited notable inhibition zones.

Molecular networking revealed a distinct cluster (**Figure 5C**) enriched in the active fractions, with several high-intensity nodes, including one at *m/z* 792.3, which was putatively annotated as the protonated adduct for TAN 1518B (*m/z* calc. 792.2862, *m/z* obsv. 792.2868, 0.8 ppm), a tetracycline-related compound, based on accurate mass (MSI Level 3). Interestingly, a second abundant feature in the same cluster (highlighted in red), *m/z* 808.3, showed a 16 Da mass shift and earlier retention time (7.5 min vs. 8.5 min), suggesting a structural modification consistent with increased polarity. While no GNPS library match was found, the spectral similarities for the UV-visible absorption patterns (**Figure 5D**) of the *m/z* 808.3 and 792.3 ions further support analogous chromophore systems and likely close structural relationships. Notably, the UV maxima and profiles resemble those reported in literature for TAN 1518B (30). Due to its higher abundance in the active fractions and spectral properties, the *m/z* 808.3 ion is a promising novel analogue of TAN 1518B. However, low yields in current culture conditions prevented its isolation and full structural elucidation. Future work will focus on optimizing production for NMR characterization.

### Co-culture-induced metabolites reveal interaction-dependent chemistry

While strain-specific specialized metabolism accounts for a significant portion of the observed chemical diversity, many bioactive features remain unexplained by monoculture analyses alone. To address this, we investigated the potential for interaction-specific metabolite production by analyzing LC-MS/MS data from the co-cultures between 36 *Pseudonocardia* strains and two *Escovopsis* isolates.

A comparative overview using a Venn diagram (**Figure 6A**) revealed a distinct subset of ions exclusively detected in co-culture conditions, absent from both monocultures and blanks according to the thresholds defined in the methodology, suggesting that microbial interactions trigger specific metabolic responses. To further investigate this interaction-dependent chemistry, we revisited the molecular network generated in **Figure 3A** and focused on clusters predominantly associated with co-cultures, particularly those involving *Pseudonocardia* and *Escovopsis* ICBG 729, because they were more abundant. Among these, two representative clusters were selected for detailed analysis based on their annotation potential (**Figure 6B–C**). These clusters included a series of lipodepsipeptides (compounds 9–10, MSI level 2) and linear peptides (compounds 11–14, MSI level 3), respectively associated with *Pseudonocardia* strains ICBG 1050 and ICBG 1288. These compounds were putatively identified as protonated adducts through GNPS spectral library matching, MS1 exact mass comparison with natural product databases, and manual inspection of fragmentation patterns (**Figures S13-19**).

**Figure 6.**
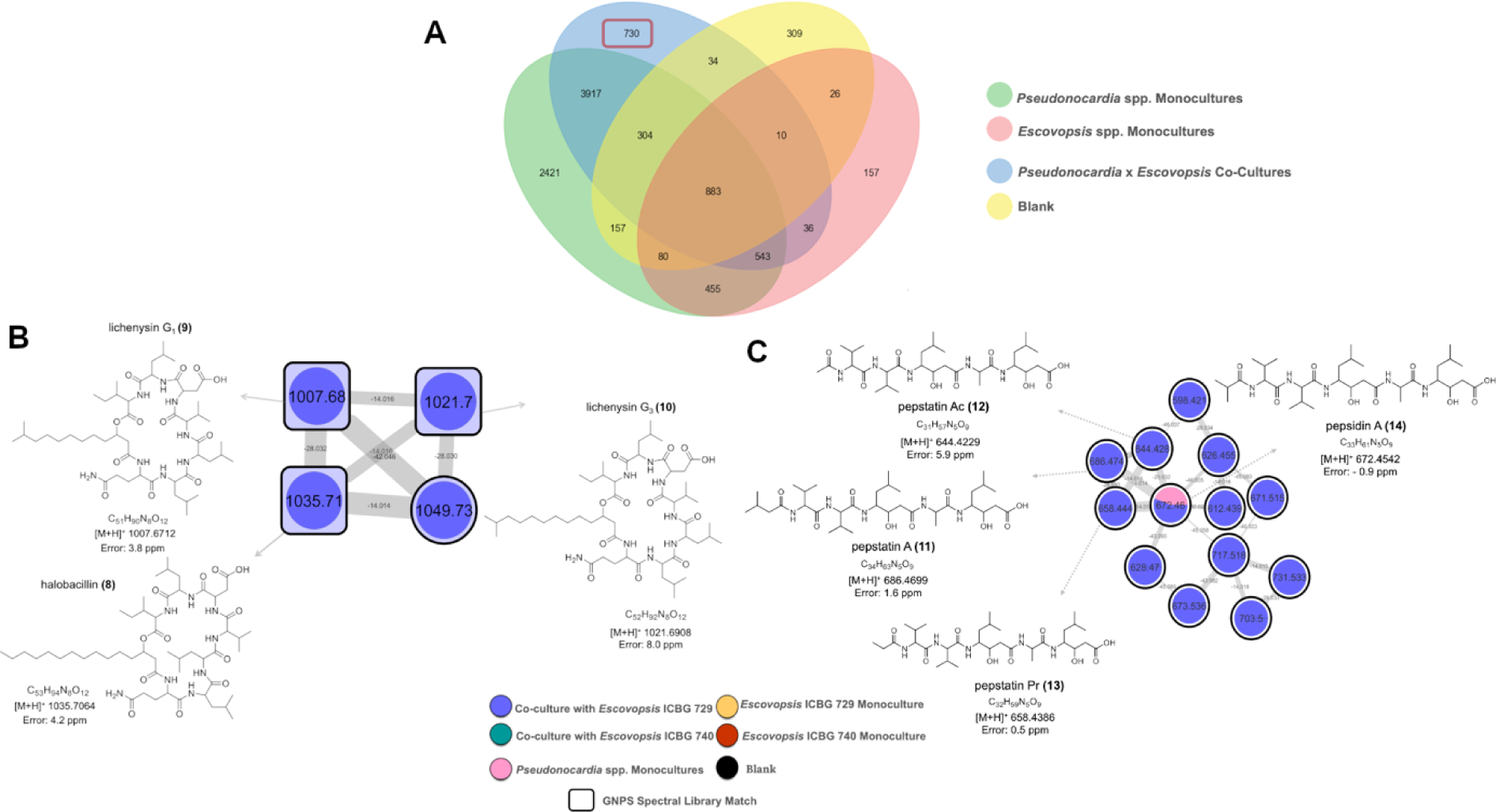
Venn diagram and molecular network clusters highlighting metabolite production in fungal-bacterial co-cultures. **(A)** Venn diagram representing ions detected by LC-MS/MS analyses of extracts from microbial interactions between 36 *Pseudonocardia* strains and two *Escovopsis* strains. Ions are classified according to their origin: *Pseudonocardia* monocultures (green), *Escovopsis* monocultures (red), co-cultures (blue), and blanks (yellow). Notably, several ions were uniquely detected in co-cultures, suggesting metabolite induction due to microbial interaction. **(B)** Molecular cluster associated with the interaction between *Pseudonocardia* ICBG 1025 and *Escovopsis* ICBG 729 and their compounds annotated via GNPS spectral library (8–10). This cluster was extracted from the molecular network generated from the LC-MS/MS data of all *Pseudonocardia–Escovopsis* interactions (Figure 3). **(C)** Molecular cluster related to the interaction between *Pseudonocardia* ICBG 1245 and *Escovopsis* ICBG 729, with compounds annotated at MSI level 3 (11–14). This cluster was also derived from the molecular network of all pairwise microbial interactions.

To further survey the metabolic capabilities of our *Pseudonocardia* collection, we also examined the distribution of attinimicin, a conserved antifungal metabolite selectively active against *Escovopsis* and previously reported across multiple regions of Brazil but absent in Panamanian strains (22). To assess whether this compound was similarly elicited under co-culture conditions, we compared its relative intensity in monocultures versus co-cultures with *Escovopsis*. Attinimicin was detected in 8 out of 36 strains (22.2%) (**Figures S20-21**). These strains were consistent with PCR-based detection of the *att* biosynthetic gene cluster (22). Interestingly, relative abundances were generally higher in co-culture than in monoculture, particularly in interactions with *Escovopsis* ICBG 740, suggesting that ecological cues enhance the production of this specialized metabolite.

In addition to the metabolites produced by *Pseudonocardia*, our dataset also revealed a suite of specialized fungal metabolites associated with *Escovopsis* spp., expanding the chemical perspective of the interaction. The Venn diagram (**Figure 6A**) already indicated the presence of ions exclusive to *Escovopsis* monocultures and ions that persisted or intensified in co-cultures, suggesting that the fungus also contributes substantially to the chemical environment during antagonistic interactions. When these ions were mapped back onto the global molecular network (**Figure 3A**), a discrete molecular cluster (**Figure 7**) emerged that contained only features associated with shearinine-type metabolites previously linked to pathogenicity and behavioral disruption in attine ants (31–34). While 22,23-dehydro shearinine A (**15**), shearinines D (**16**) and F (**17**) were annotated using the GNPS library comparison (MSI level 2), shearinines B (**18**), E (**19**), G (**20**) and H (**21**) were attributed through MS1 comparison with public natural products databases (MSI level 3) (**Figures S22-24**). According to the colors of the pie chart, it is possible to observe that the detection of these analogs presented diverse patterns varying between the two fungal strains (ICBG 729 and ICBG 740) and between monoculture and co-culture conditions. Notably, shearinine F was the only compound consistently detected across all conditions, whereas shearinine B was restricted to *Escovopsis* ICBG 729 monocultures, and 22,23-dehydro-shearinine A, shearinine E, shearinine G, and shearinine H were enriched in ICBG 740.

**Figure 7.**
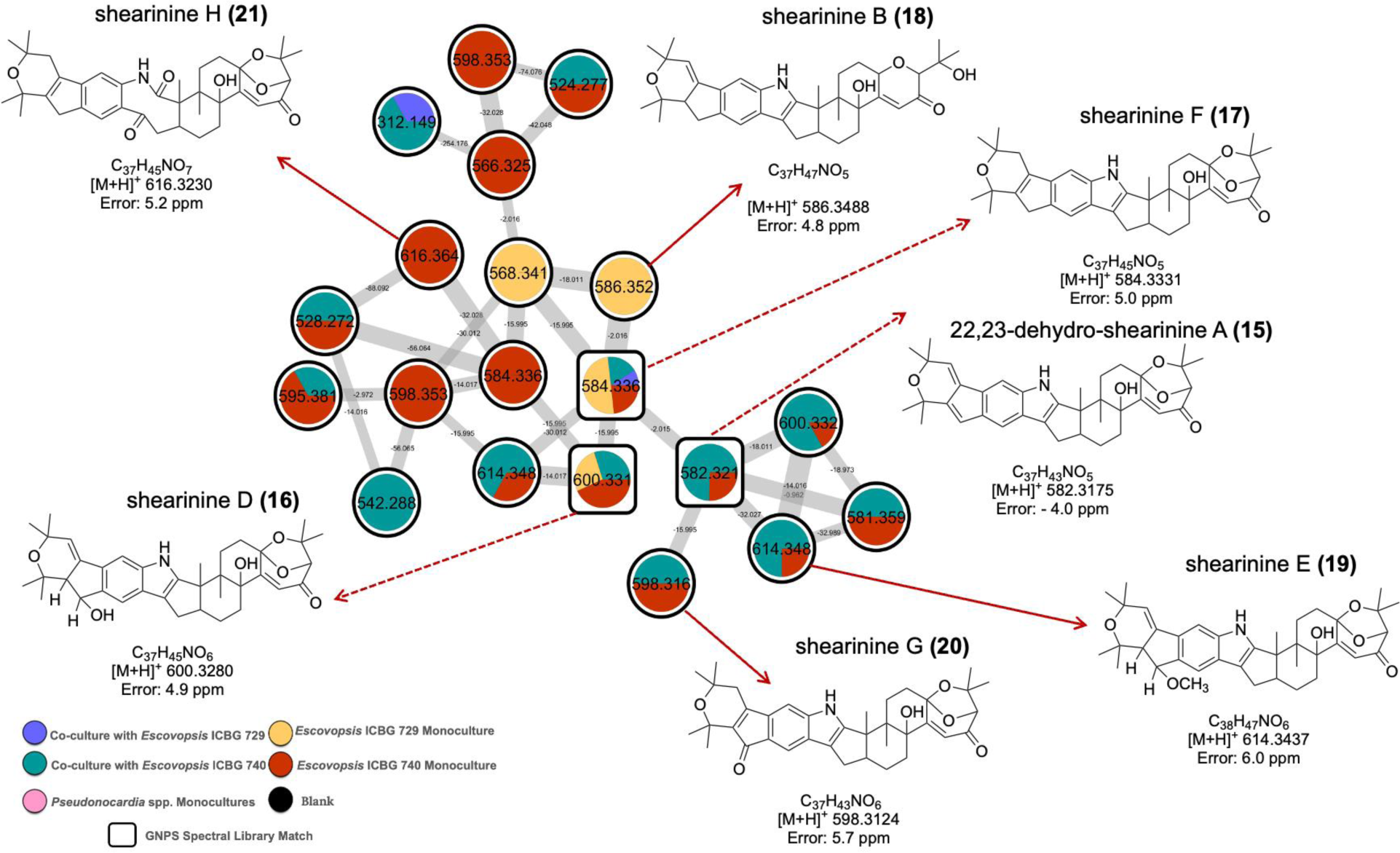
Molecular cluster associated with the interaction between 36 *Pseudonocardia* strains and *Escovopsis* ICBG 729 and ICBG 740. Cluster extracted from the global molecular network (Figure 3) showing ions annotated as the protonated adducts of shearinine-type metabolites via GNPS spectral library matches and MS1 exact-mass comparison with public natural product databases. Node colors correspond to sample types as indicated in the legend, and pie charts indicate the relative abundance of each feature in the different culture condition. Black nodes represent blank samples. Nodes outlined in white indicate GNPS library matches. The chemical structures correspond to the annotated analogs: 22,23-dehydro-shearinine A, shearinines B, D, E, F, G, and H.

## DISCUSSION

The antagonism assays revealed striking variability in the inhibitory potential of *Pseudonocardia* symbionts against *Escovopsis*, underscoring the functional diversity that exists even among closely related strains. While some isolates displayed little to no suppression of fungal growth, others achieved complete inhibition, highlighting a broad spectrum of defensive capacities. Notably, inhibition was consistently stronger and more variable against *Escovopsis* ICBG 729, which was isolated from the same ant genus (*Paratrachymyrmex*) as the bacterial strains, compared to ICBG 740 from *Acromyrmex* colony. This pattern suggests that the effectiveness of symbiont defense may be enhanced when bacteria and pathogens share a common evolutionary and ecological context, pointing to the importance of local adaptation and co-evolutionary history in shaping the outcomes of these multipartite interactions. These differences in the functional results emphasize that the defensive roles of Amazonian *Pseudonocardia* strains are not uniform but instead reflects strain-specific ecological matching between bacterial symbionts and their fungal antagonists, consistent with previous studies showing host-specificity and co-evolutionary dynamics between *Pseudonocardia*, *Escovopsis*, and their ant hosts (7, 19, 35, 36). In addition, given the extensive environmental heterogeneity of the Amazon biome, from soil type and humidity to microbial and fungal community composition, it is plausible that *Pseudonocardia* symbionts have evolved distinct defensive responses under localized selective pressures, further shaping the variability observed in our antagonism assays (37).

The functional variability in inhibition aligns with our metabolomic analyses, which revealed that *Pseudonocardia* strains harbor highly diversified and strain-specific chemical repertoires that do not strictly follow phylogenetic relatedness. A similar decoupling between taxonomy and metabolite output has been described in other actinobacteria, including *Streptomyces*, where closely related strains can display substantially different secondary metabolomes (38–40). This diversification may reflect selective pressures imposed not only by fungal antagonists such as *Escovopsis* but also by competition among *Pseudonocardia* themselves. Vertically transmitted strains have been shown to produce antibiotics active against other actinomycetes, ensuring dominance within the colony, while horizontal gene transfer further contributes to biosynthetic diversification (41–45). Interestingly, these patterns contrast with recent work on free-living *Pseudonocardia*, where biosynthetic and metabolomic repertoires closely mirror phylogenetic relationships (46). The divergence observed in ant-associated strains suggests that symbiotic integration imposes unique ecological pressures that decouple metabolic output from taxonomy and drive diversification of specialized metabolites.

To further explore strain-level metabolic repertoire diversity, we selected three *Pseudonocardia* isolates (ICBG 1122, ICBG 1025, and ICBG 1860) that exhibited distinct chemical and phenotypic signatures. Despite their close phylogenetic 16 S relatedness, these strains displayed markedly divergent metabolomic profiles, reinforcing the idea that ecological pressures and horizontal gene transfer, rather than taxonomy alone, underpin biosynthetic diversification. In particular, the pronounced chemical divergence observed in strains ICBG 1122 and ICBG 1860 raises the possibility that accessory genetic elements, such as strain-specific plasmids or horizontally acquired biosynthetic gene clusters, may contribute to their distinctive metabolomic and antagonistic phenotypes. Such elements, which are not captured by 16S rRNA gene analysis, are known to play an important role in shaping secondary metabolite production in actinobacteria through horizontal gene transfer and genome plasticity (45, 47).

Strain ICBG 1122 produced dentigerumycin analogs, including dentigerumycin F, a member of a cyclic depsipeptide family originally described from *Apterostigma dentigerum*-associated *Pseudonocardia* with selective activity against *Escovopsis*. In contrast, ICBG 1025 yielded provipeptide A and the β-carbolines norharman and harman. β-carbolines are indole alkaloids broadly distributed across plants, fungi, tunicates, and actinobacteria, and have been implicated both as signaling molecules that induce secondary metabolism in *Streptomyces* and as antimicrobials against diverse pathogens (48–53). Provipeptide A, originally isolated from marine *Streptomyces associated* with *Bryopsis* algae, exhibits antibacterial activity against phytopathogens and human pathogens, including *S. aureus* (29). Meanwhile, *Pseudonocardia* ICBG 1860 was enriched in tetracycline-related compounds, including TAN 1518B, a known DNA topoisomerase I inhibitor with antitumoral activity, along with a putative novel analog (30). These findings exemplify the chemical richness harbored by individual strains, with implications for natural product discovery, and shows the untapped biosynthetic potential of symbiotic actinomycetes from understudied regions such as the Amazon. Furthermore, our methodological approach proved effective in linking chemical diversity to bioactivity. By prioritizing fractions of crude extracts through molecular networking, we were able to target those directly associated with inhibitory activity against *Escovopsis* and human pathogens, validating the power of metabolomic-guided chemical screening as a strategy to identify functionally relevant metabolites. Importantly, the biological activities previously reported for these metabolite classes may partly account for the inhibitory effects observed in our bioassays, reinforcing the connection between the chemical profiles detected and the antimicrobial phenotypes evaluated.

Beyond strain-specific repertoires, co-culture experiments revealed context-dependent activation of biosynthetic pathways, with metabolites emerging exclusively in the presence of *Escovopsis*. These findings reveal that interspecies interactions act as ecological triggers, likely mimicking natural cues within the ant colony environment (15, 21, 54). The recurrent induction of metabolites under co-culture conditions was readily visualized through Venn diagram comparisons and molecular networking of LC-MS/MS data. Notably, hallobacillins, lichenisins, and pepstatins were annotated in molecular clusters exclusively associated with *Pseudonocardia*–*Escovopsis* interactions. Given their bacterial origin and reported antifungal properties, these compounds are likely produced by *Pseudonocardia* in response to *Escovopsis*, partly explaining the inhibitory activity observed in our bioassays (55–57). In addition to these newly annotated clusters, we also examined the conserved antifungal metabolite attinimicin, previously reported across Brazil but absent in Panamanian strains (22). It was detected in 8 out of 36 strains (22.2%), with higher relative abundances under co-culture conditions, particularly with *Escovopsis* ICBG 740, suggesting that ecological cues enhance its production. These results align with recent advances in co-culture studies between pathogenic *Escovopsis* and symbiotic *Pseudonocardia* in the attine ant system. Gemperline et al. localized pathogen-induced *Pseudonocardia* metabolites on the ant cuticle via mass spectrometry imaging, providing direct spatial evidence of chemical defense (20). Boya et al. used imaging and molecular networking to map the antagonistic interface between *Streptomyces* and *Escovopsis*, though outside a symbiotic context (58). Most notably, it has been previously demonstrated that the biosynthesis of chemical defenses in both *Pseudonocardia* ICBG 1122 and *Escovopsis* ICBG 729 can be elicited by exposing each microorganism to the supernatant containing secondary metabolites produced by its competitor (21).

The detection of shearinines is consistent with prior studies demonstrating that these molecules are major pathogenicity factors produced by *Escovopsis*, capable of killing *Pseudonocardia* symbionts and altering the behavior and ultimately the survival of ant workers (31, 32, 34). The presence of multiple shearinine analogs in our Amazonian isolates, including analogs not previously described for ant-associated *Escovopsis* (59), reinforces the idea that the fungus possesses a potent and chemically diverse offensive arsenal. Moreover, the overlap between fungal-specific ions and co-culture ions in the Venn diagram suggests that *Escovopsis* upregulates at least part of this arsenal in response to bacterial competition, consistent with ecological observations of pathogen-induced garden collapse in attine ant colonies. In this case, these results reveal that antagonistic interactions within the *Pseudonocardia*–*Escovopsis* system are chemically bidirectional and that *Escovopsis* also contributes with a rich repertoire of specialized metabolites that likely shape the outcome of the symbiosis.

## CONCLUSION

Our findings demonstrate the potential of microbial interactions to activate silent biosynthetic pathways in *Pseudonocardia* symbionts while also revealing important gaps in understanding their chemical diversity. Even 16 S phylogenetically closely related strains displayed distinct metabolomic repertoires and interaction-dependent chemistries, and functional outcomes against *Escovopsis* directly reflected this underlying metabolomic diversification. This contrasts with free-living *Pseudonocardia*, where specialized metabolism more closely mirrors phylogeny. By systematically characterizing multiple Amazonian strains, we showed that defensive roles in these symbioses are versatile, strain-specific and chemically bidirectional. However, the lack of genome-resolved sequencing currently limits our ability to distinguish the relative contributions of core and accessory biosynthetic gene clusters to this metabolic diversification. Future integrative studies coupling metabolomics with genomic and transcriptomic data will be key to unraveling the functional and evolutionary basis of this remarkable metabolic variability.

## MATERIAL AND METHODS

### General experimental procedures

Ultra high-performance liquid chromatography-high resolution electrospray-ionization mass spectrometry (UHPLC-HR-ESI-MS) data were obtained at the Collaborative Mass Spectrometry Innovation Center da University of California - San Diego (UCSD) using an UHPLC Ultimate 3000 (Thermo Scientific) system equipped with a Maxis Q-TOF (Bruker Daltonics) mass spectrometer. HPLC purifications were carried out using a semi-preparative system (Shimadzu Nexera) equipped with a photodiode array detector. NMR spectra were recorded using a Bruker Avance® DRX-600 MHz spectrometer equipped with a helium High Sensitivity Prodigy Cryoprobe (600 and 150 MHz for ^1^H and ^13^C NMR, respectively) at Department of Chemistry, Federal University of São Carlos (UFSCar).

### Microbial Strains

For this study, we selected 36 actinobacterial strains belonging to the genus *Pseudonocardia*, isolated from *Paratrachymyrmex* ants collected in the Amazon region (Anavilhanas National Park and Adolfo Ducke Forest Reserve). Additionally, two *Escovopsis* strains were selected, isolated from *Paratrachymyrmex* and *Acromyrmex* ants collected in the Anavilhanas National Park. The identification of *Pseudonocardia* and *Escovopsis* strains were previously reported (21, 22). Identification codes from the International Cooperative Biodiversity Groups (ICBG) project and the geographic coordinates for the *Pseudonocardia* and *Escovopsis* strains are provided in the Supplementary Material. Legal permits for the collection of biological samples and access to Brazilian genetic material were granted by SISBIO (46555–8) and SISGEN (A25AA57; A9D808C), respectively.

### Microbial Cultivation

*Pseudonocardia* strains were grown on ISP-2 agar (0.4% glucose, 1% malt extract, 0.4% yeast extract, 0.2% agar) and *Escovopsis* strains on PDA (0.4% potato, 2% dextrose, 1.5% agar) in 60 mm Petri dishes (25 mL) at 30 °C for 30 days. Spores were collected in sterile 20% glycerol, stored at -80 °C, and used to inoculate interspecies interaction assays. Monocultures were prepared by spotting 100 µL of spore suspension at one plate delimited edge and incubating for 14 days at 30 °C. Co-cultures were initiated by inoculating *Pseudonocardia* as above, adding the same amount of *Escovopsis* inoculum on the opposite delimited edge after 14 days, and incubating for additional 14 days at 30 °C before extraction.

Small scale cultivations were made to confirm the presence of differential metabolites observed in metabolomic analyses and correlate them with bioactivities, particularly for strains ICBG 1122, ICBG 1025, and ICBG 1860. For each strain, 100 µL of pre-culture was inoculated into five medium-sized Petri dishes (90 × 15 mm) containing 30 mL of ISP-2 agar and incubated at 30 °C for 14 days prior to extraction. Subsequently, the same strains were cultivated on a larger scale using 150 plates for ICBG 1122, 140 plates for ICBG 1025, and 100 plates for ICBG 1860 (150 × 15 mm, each containing 70 mL ISP-2 agar). Plates were inoculated with 500 µL of pure pre-culture in ISP-2 medium and incubated at 30 °C for 14 days before extraction.

### 16S rRNA Gene-Based Phylogenetic Analysis

Phylogenetic reconstruction was performed using 16S rRNA gene sequences obtained from previous studies conducted by our research group, generated using Illumina MiSeq and PacBio sequencing platforms (22). Sequence alignment and phylogenetic analyses were conducted in MEGA v7.0.26 (60). Phylogenetic relationships were inferred using the maximum likelihood method under the general time-reversible (GTR) nucleotide substitution model with a discrete gamma distribution to account for rate heterogeneity among sites (61). The analysis included 36 nucleotide sequences, and all positions containing gaps or missing data were excluded. The final aligned dataset comprised 3,105 nucleotide positions. Branch support was assessed using 1,000 bootstrap pseudoreplicates. The model allowed for a proportion of evolutionarily invariant sites ([+I], 39.52%). Phylogenetic trees were visualized and edited using iTOL v6 (https://itol.embl.de/) (62).

### Metabolite Extraction and LC-MS/MS Sample Preparation

Cultures for metabolomic analysis were immersed in ethyl acetate and extracted for 24 h at room temperature, and the organic phase was filtered and evaporated at 40 °C under reduced pressure to yield crude extracts. These extracts were fractionated using C18 solid-phase extraction (100 mg, 1 mL; Supelco), preconditioned with MeOH and water, washed with water, and eluted with 90% aqueous MeOH. The same metabolite extraction protocol was employed for small and larger scales cultures, but different C18 cartridges were used for fractionation (500 mg/5 mL and 10 g/100 mL respectively). Eluates were dried at 40 °C under reduced pressure, resuspended in MeOH to reach 1 mg/mL, sonicated for 15 min, and centrifuged at 12,000 rpm for 15 min. For metabolomic samples MeOH containing sulfamethazine was added at 2 μmol/L.

### LC-MS/MS Analysis

A C_18_ Phenomenex Kinetex 100 Å 50 x 2.1 mm UPLC column with particle size 1.7 µm was used on a Bruker Elute UPLC. The column was equilibrated to 95% A (H_2_O + 0.1% formic acid) and subjected to a 10 min gradient from 5 to 100% B (ACN + 0.1% formic acid) with an injection volume of 10 μL and a flow rate of 0.5 mL/min at 40 °C. Data was acquired in Compass Hystar 6.2 software. The high resolution LC-MS/MS detection were performed on in positive mode from *m*/*z* 50–2000. Nebulizer gas (N_2_) was set to 2 bar, and dry gas was set to 9.0 L/min. Dry temperature was set to 200 °C. The ESI conditions were set with the capillary voltage at 4.5 kV. LC-MS/MS data was collected using data-dependent acquisition (DDA) mode at an MS spectra rate of 3 Hz and MS/MS spectra rate at 10 Hz with the top 5 precursors selected for fragmentation at the collision energies and isolation widths shown below.

| Mass (*m/z*) | Width (*m/z*) | Collision Energy (eV) |
| --- | --- | --- |
| 50 | 2 | 20 |
| 1000 | 4 | 20 |
| 500 | 6 | 20 |
| 1300 | 8 | 30 |

### LC-MS Data Processing

To create molecular networks in the GNPS environment, the tandem mass spectra data were converted into .mzXML files using MSConvert (63)and submitted to the GNPS platform (25) with the following parameters: all MS/MS peaks within ±17 Da of the precursor *m/z* were removed. Repeated spectra were grouped using the MS-Cluster algorithm (Classic Molecular Networking) with a precursor ion mass tolerance and fragment ion tolerance of 0.02 Da to generate consensus spectra, each represented as a node. Only consensus spectra containing at least two nearly identical spectra were considered. A molecular network was then constructed using the representative consensus MS/MS spectra, with edges filtered to retain only those with a cosine similarity score above 0.7 and at least four matching fragment peaks. Edges were maintained in the network only if each node appeared within the 10 most similar nodes of the other. The network spectra were subsequently matched against GNPS spectral libraries, which were filtered in the same manner as the input data. Library matches were retained if they had a cosine score above 0.7 and at least four matching peaks. The resulting output was imported and visualized as molecular networks in Cytoscape 3.4.0 (64).

A featured list was created with the raw MS1 Bruker files, processed using T-ReX® 3D in MetaboScape (Bruker Daltonik) and used to perform the multivariate analysis. The minimum number of features for extraction and result was 3/18, so a feature must have been present in 3 files to appear in the feature list. The peak intensity threshold was 2500, retention time spanning the gradient only. Ions used for extraction include [M + H]^+^, [M + Na]^+^, [M + NH_4_]^+^, and [M + K]^+^. Modifications included a loss of water [M – H_2_O + H]^+^. Ion intensities were normalized using Total Ion Current (TIC) normalization to account for technical variation between runs. Following feature list generation, graphic representations including the heat map, Venn diagram and Procrustes analysis were made and visualized in Python using the scikit-learn, matplotlib, and seaborn packages within a Jupyter Notebook, an open-source, web-based interactive computing environment.

For the heat map, similarities in the metabolome relative intensities detection among strains were calculated using Euclidean distance and visualized by hierarchical clustering heat maps generated using the Seaborn package. Grouping and overlap of metabolite features among strain sets were visualized using Venn diagrams generated with the Matplotlib package.

Procrustes analysis was performed to compare ordinations derived from 16S rRNA gene–based phylogenetic distances and metabolite feature abundance data. Bray–Curtis dissimilarity matrices were generated for both datasets in QIIME v1.9.1, followed by independent Principal Coordinates Analysis (PCoA). Ordinations were aligned using Procrustes analysis in QIIME v1.9.1, and visualized using EMPeror (65).

GNPS molecular networking jobs are deposited under the following task IDs: https://gnps.ucsd.edu/ProteoSAFe/status.jsp?task=317e5abbc0674930bde85e83afb26b71 https://gnps.ucsd.edu/ProteoSAFe/status.jsp?task=35fc19929cc640178255fde075cb828b

### Antimicrobial Bioassays of Fractions from Pseudonocardia Extracts

*Pseudomonas aeruginosa* ATCC 15442, *Candida albicans* INCA40006, and *Staphylococcus aureus* INCA00039 were grown in liquid ISP-2 at 30 °C, 170 rpm for 24 h. Pour plate assays were prepared by mixing the individual cultures with ISP-2 soft agar (0.7% agar) to an OD₆ ₀ ₀ of 0.1, pouring 20 mL into 150 × 15 mm Petri dishes, and applying 10 μL of extracts fractionated (10 mg/mL in DMSO or MeOH:H₂ O 1:1) at 2 cm spacing. Plates were incubated at 30 °C, and inhibition halos were recorded after 24 h. For *Escovopsis*, ISP-2 soft agar containing 10⁵ spores/mL was poured over into 150 × 15 mm Petri dishes, followed by extract application as above. Inhibition halos were assessed after 7 and 14 days at 30 °C.

### Metabolites Isolation

The resulting MeOH fractions from large-scale cultivation of *Pseudonocardia* strains ICBG 1122 and ICBG 1025 were dried under reduced pressure and subjected to further analysis. For strain ICBG 1122, the 75% and 100% MeOH fractions afforded 2.3 mg of dentigerumycin F. For strain ICBG 1025, the 50% and 75% MeOH fractions yielded 0.65 mg of provipeptide A. Both compounds were isolated by semi-preparative HPLC using reversed-phase C18 (Phenomenex Gemini, C18, 5 μm, 250 × 10 mm, 35 °C) and C6-phenyl columns (Phenomenex Gemini, C6-phenyl, 5 μm, 250 × 10 mm, 35 °C). Samples were prepared at 3 mg/mL in ACN:H₂ O (1:1) and injected at 200 μL. Detection was carried out by DAD monitoring at 190 and 254 nm. For dentigerumycin F (ICBG 1122, C18 column), the gradient was 60% ACN for 5 min, ramped to 100% over 35 min, and held at 100% for 10 min. For provipeptide A (ICBG 1025, C6-phenyl column), the gradient was 10% ACN for 10 min, ramped to 50% over 30 min, held for 10 min, then ramped to 100% over 10 min and held at 100% for 5 min.

### antiSMASH BGC Annotation

The putative biosynthetic gene cluster for dentigerumycin-like compounds were annotated from a previously sequenced genome of *Pseudonocardia* sp. ICBG 1122 using antiSMASH v7.0 with default settings, including cluster detection for polyketide synthases (PKS), nonribosomal peptide synthetases (NRPS), hybrid clusters, and other secondary metabolite biosynthetic pathways (22, 66). Selection criteria included a minimum similarity of ≥50% to a known BGC associated with a characterized compound. Regions located at sequence contig edges were excluded to avoid incomplete or ambiguous annotations.

## ACKNOWLEDGEMENTS

This research was supported by the São Paulo Research Foundation (FAPESP) grants #2013/50954-0 (MTP), #2013/07600-3 (MTP; CEPID-CIBFar), #2020/06430-0 (COG), #2019/06061-8 (COG), #2015/01001-6 (WGPM), #2016/17614-0 (WGPM); #2025/03943-0 and #2020/02207-5 (RRDS); by the Conselho Nacional de Pesquisa e Desenvolvimento Tecnológico (CNPq) grants #303792/2018-2 and #307893/2022-7 (MTP); and by the Coordenação de Aperfeiçoamento de Pessoal de Nível Superior - Brasil (CAPES) - Finance Code 001. AMC-R and PCD were supported by the Gordon and Betty Moore Foundation, GBMF12120 and https://doi.org/10.37807/GBMF12120.

